# Running rabid: modelling of European bat lyssavirus (EBLV) pathways in a bat population

**DOI:** 10.1101/795856

**Authors:** Carys Breeze, James Aegerter, Graham C Smith

## Abstract

Worldwide the 16 species of lyssaviruses all exhibit a similar pathology in most mammals, including man; with successful infections usually ending with death. Recently it has been demonstrated that European bat lyssaviruses (EBLV) are not invariably fatal in their wild reservoir host bat species, however the mechanisms and epidemiological consequences of this resistance are interesting and unexplored and the fundamental pathology in bats is still unclear. Here we modelled alternative pathological pathways to explore which appear most plausible, with respect to our limited knowledge of bat-rabies epidemiology and also host population dynamics. Two models were created, one based on a standard progression of disease (classic SEIR model) and the other modified to allow for animals to become either rabid or immune (flexible model). Of these our flexible model was found to be more plausible, demonstrating a much lower sensitivity to epidemiological parameters and by inference the more likely to represent the real-life process occurring in wild European bat populations, with a comparative state space ratio of 1:47. This result implies that it is highly probable survival and post-infection immunity is a widespread epidemiological phenomenon rather than an infrequent consequence of an aborted infection in few individuals. These results can be used to inform laboratory studies on bat immunology and future bat modelling work.

## 1. Introduction

There are currently 16 recognised Lyssavirus species, with more awaiting classification (Maes et al., 2019); the most widespread and well-known of these is classical rabies (RABV). Almost all are recorded as producing a similar pathology in mammalian hosts, with a successful muscle infection leading to invasion of the nervous system, usually followed by death in untreated cases (Lafon, 2005), with very few documented cases of survival in humans, even after radical intervention (e.g. de Souza & Madhusudana, 2014), or in other animals, although both aborted infection and apparent recovery have been documented (Fekadu, 1991). Whilst RABV has been shown to circulate in a number of mammalian families worldwide (e.g. Canidae, Procyonidae) it has also been associated with New World bats (e.g. Vespertilionidae, Phyllostomidae) such as the common vampire (*Desmodus rotundus*) (Baer, 1991) most other Lyssaviruses are associated with a variety of bat hosts (Kuzmin & Rupprecht, 2015). Australian bat Lyssavirus (ABLV) is commonly reported in fruit bats (*Pteropus* spp.) and an insectivorous bat (*Saccolaimus aliventris*); in Eurasia, Aravan virus (ARAV) and Khujand virus (KHUV) are associated with *Myotis* bats and West Caucasian bat virus (WCBV) again associated with *Mi. schreibersii* (Banyard et al., 2011; Kuzmin & Rupprecht, 2015). The rabies strains considered present in European bats are European bat lyssavirus (EBLV) types 1 and 2 and Bokeloh virus (BBLV). EBLV-1 is typically associated with Serotine bats (*Eptesicus sp.*) and EBLV-2 has been recorded in two species of *Myotis* bat (*M*. daubentonii & *M*. dasycneme) (Banyard et al., 2011), BBLV in *M*. nattereri and Llieda bat virus associated with the bent-winged bat (*Miniopterus schreibersii*). Instances of EBLV spill-over have been found in a stone marten in Germany, and a cat and two sheep in Denmark (Harris et al., 2006). Four people have known to have contracted rabies infection due to contact with EBLV in Europe (Fooks et al., 2003).

Unlike other unvaccinated mammals, bats may produce and maintain detectable titres of rabies neutralising antibodies with no signs of disease (Arguin et al., 2002; Bowen et al., 2013; Harris et al., 2009; Rønsholt et al., 1998), and may survive with antibodies for many years (Amengual et al., 2008; Serra-Cobo et al., 2002); further, some individuals can survive experimental challenge (Almeida et al., 2005; Freuling et al., 2009; Hughes et al., 2006; Jackson et al., 2008). It has been speculated that their survival of a rabies challenge may be related to their unusual lifestyles or life history (Luis et al., 2013; O’Shea et al., 2014), or details of their immune system (Zhou et al., 2016).

There are two potential routes to achieving immunity. In most mammals untreated symptomatic rabies is understood to be invariably fatal and antibody production is usually associated with the terminal phase of symptoms including virus secretion (infectiousness), so the presence of antibodies may indicate survival from symptomatic rabies (i.e. recovery from disease or post-infectious immunity), which has been reported rarely (Fekadu & Baer, 1980). However, sometimes healthy unvaccinated animals (Fekadu, 1991) and people (Gilbert et al., 2012) have been detected with rabies neutralising antibodies, suggestive of an acquired immunity following exposure (e.g. aborted infection, cleared sub-clinical infection, cleared asymptomatic disease) which we interpret here as immunity without a period of infectiousness. We will refer to these two options as a classical post-infectious immunity and a flexible pre-infectious immunity, and recognise any population may be composed of individuals capable of following either routes.

Here, we create two alternative models of rabies in a bat population to explore these competing mechanisms. We explore the population level responses of these two processes and estimate the possible variance in the input parameters that will lead to plausible epidemiological behaviour (e.g. realistic population levels of seroprevalence, population decline and number of active rabies cases). We then quantify the parameter space supporting plausible epidemiological observations and identify the most likely model pathology which describes this. We argue that if one of these models has a restricted input parameter space compared to the other, then the restricted model is less likely.

## 2. Methods

### 2.1 Disease models

Two different models were produced in the modelling and simulation programme STELLA^®^ (version 9.1: iSee Systems Inc., Lebanon, NH) to represent realistic bat host population dynamics whilst also depicting the alternative epidemiologies. Both models were based on a typical *S*usceptible-*E*xposed-*I*nfectious-*R*ecovered approach and included functions to ensure realistic disease die out if the models predicted very low levels of infectious individuals (fractions of one animal).

The classic post-infectious immunity model (equations 1-4), considers the exposure of susceptible (*S*) bats to a lyssavirus proceeding to an infected though un-infectious state (*E*). Infected animals can then become diseased (i.e. symptomatically rabid and infectious; *I*). Individuals then either die from rabies or recover and reach an immune and antibody positive state.

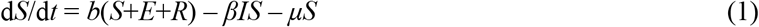

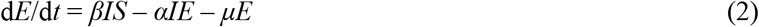

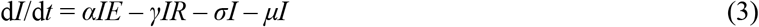

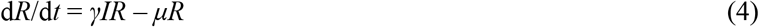

Therefore the total population is described by:

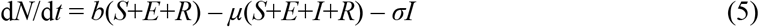

Where: *α* = rate of becoming infectious, *β* = infection rate, *γ* = rate of becoming immune, *μ* = death rate, *σ* = disease death rate, and *b* = birth rate.

In our flexible pre-infectious model, exposure of susceptible bats proceeds to them becoming infected (*E*) (eq 6). However, infected animals (eq 7) can then either become diseased (*I*) and subjected to increased mortality (eq 8), or alternatively, proceed to directly to an immune state where the animals are assumed to be producing antibodies (eq 9) where the total population is still described by eq. 5.

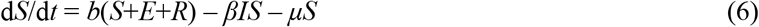

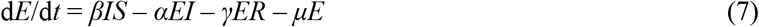

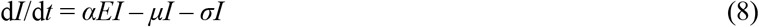

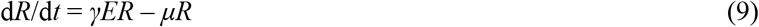

Both models share parameters and an underpinning host population dynamic in the absence of disease (Table 1). Bats have a strongly seasonal biology, so graphical functions in STELLA^®^ were used to permit seasonal fluctuations in birth and death rates. Birth rate was taken from a study on brown long-eared bats (Boyd & Stebbings, 1989) corrected for simulating only the female population. Births peak in June, but since these young remain dependent on their mother for a period of time, the overall death rate was increased over summer to allow for the death of dependent pups (Dietz & Kalko, 2006). Hibernation was not considered to affect mortality (Sendor & Simon, 2003), so this was set as a monthly constant. The epidemiological consequences of hibernation interrupting disease progression have been studied in raccoon-dogs suggesting an almost complete stasis in individual pathology with the disease only progressing upon the animal awakening (Singer et al., 2009). As the northern temperate bat species considered here all require the use of successive bouts of prolonged deep torpor to survive the late/autumn/winter/early spring period which we suggest can lead to similar effects as those seen in true hibernating animals (George et al., 2011), explaining our assumption of reduced winter disease-induced mortality.

**Table 1:**
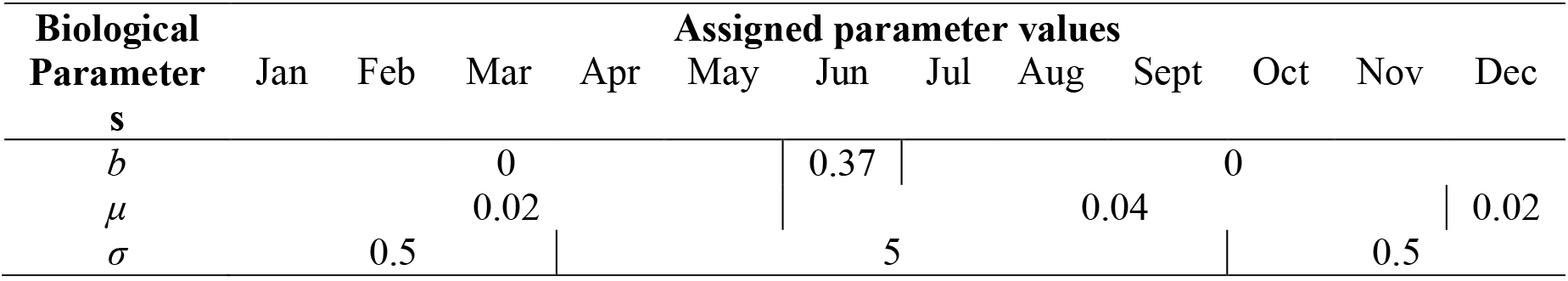
Seasonal parameters used for both models.

Initial values for disease parameters were obtained from the literature (Table 2) as a starting point and then adjusted to ensure a stable model output. Of necessity, the resultant values were slightly different for each model. Plausible epidemiological outcomes were defined to consistently evaluate alternative models and parameters. Detailed robust fine-scaled descriptions of bat population dynamics are not available, requiring the use of national scale populations trends (Singer et al., 2009), with recent stable or increasing descriptions of the Daubenton’s bat in the UK (the host of EBLV-2), though with wide confidence limits (Barlow et al., 2015). Thus we defined that a plausible output population must not decline by more than 33%. Plausible limits of rabies anti-body seroprevalence were defined as varying between 1-10%. Observational studies of classical rabies in big brown bats (*Eptiscus fucus*) recorded seroprevalence rates varied from 2-23% (Shankar et al., 2004), while rates of antibody seroprevelence for EBLV-2 in Daubenton’s bats varied from 1-5% (Harris et al., 2006). With only five recorded EBLV-2 positive Daubenton’s bats in the UK between 2006 and 2010 (Schatz et al., 2013) we set an upper limit of 1% of the population being rabid (i.e. symptomatic disease).

**Table 2.**
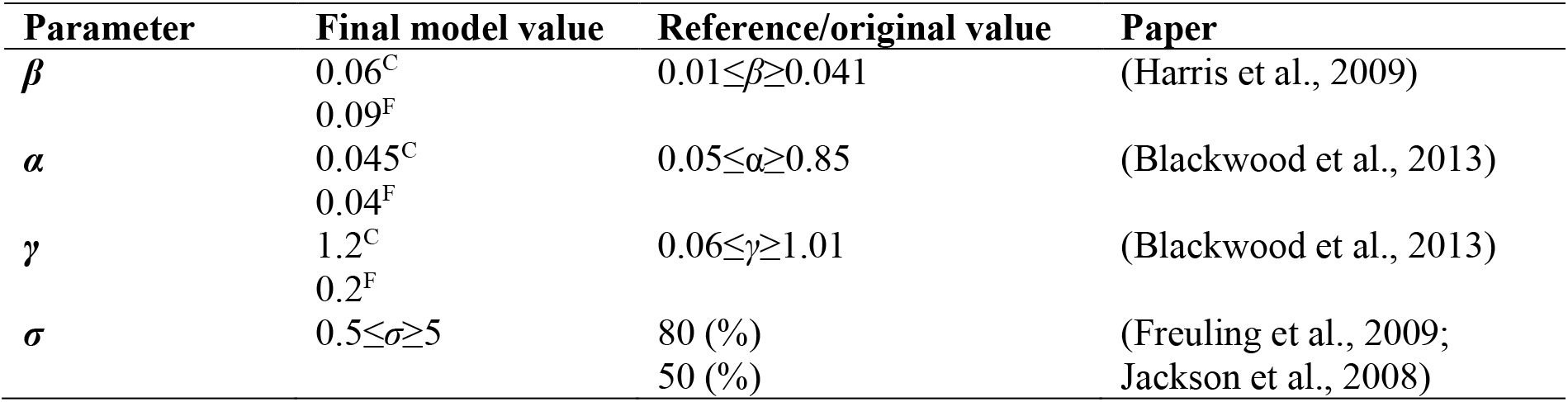
List of the disease parameter values used for the models. Where: β = infection rate, α = rate of becoming infectious, γ = rate of becoming immune, and σ = disease death rate. Where values differ between models this is indicated by C for the classical or F for the flexible model.

### 2.2 Sensitivity analysis

We performed a full sensitivity analysis to determine the limit of each input value (*β, α*, and *γ*) which produced plausible epidemiological outcomes, assuming animals in the immune state were equivalent to seropositive animals. The full input state space volume for each model was then calculated across the limiting bounds for each parameter. As we are attempting to explore real epidemiological behaviour in the wild, towards the edge of bat host species’ range and in an extremely unpredictable oceanic climate we use the following logic. There is no evidence of substantial changes in measures of either host populations or epidemiological markers of disease despite an assumption of annual variation in the dynamics of both, driven by substantial annual differences in weather and the experience and behaviour of the bat hosts. The long-term maintenance of a plausible epidemiological model requiring a narrow state-space seems less likely than one which demonstrates a more flexible and open parameter space. Narrow and specific requirements for epidemiological maintenance within a dynamic forcing environment are likely to promote unstable disease dynamics leading to either fade-out or burn-out. Thus we assume that the model with the larger input state space was more likely to reflect real-life epidemiology. We accept that this is not a form of proof, but rather allows less plausible hypotheses to be rejected and can be used to direct future studies on bat immunology.

## 3. Results

### 3.1 Classic SEIR Model - post-infectious immunity

Output for the classic SEIR model with the initial parameter values indicated a very long-term damped oscillation on top of the annual cycles (Figure 1). After an initial seropositive peak at 6%, this declined to stabilise around 2-3%. The percentage of rabid animals present in the system shows a maximum of approximately 0.3% before dropping down to around 0.05% and the population size stabilised at around 80-90%.

**Figure 1.**
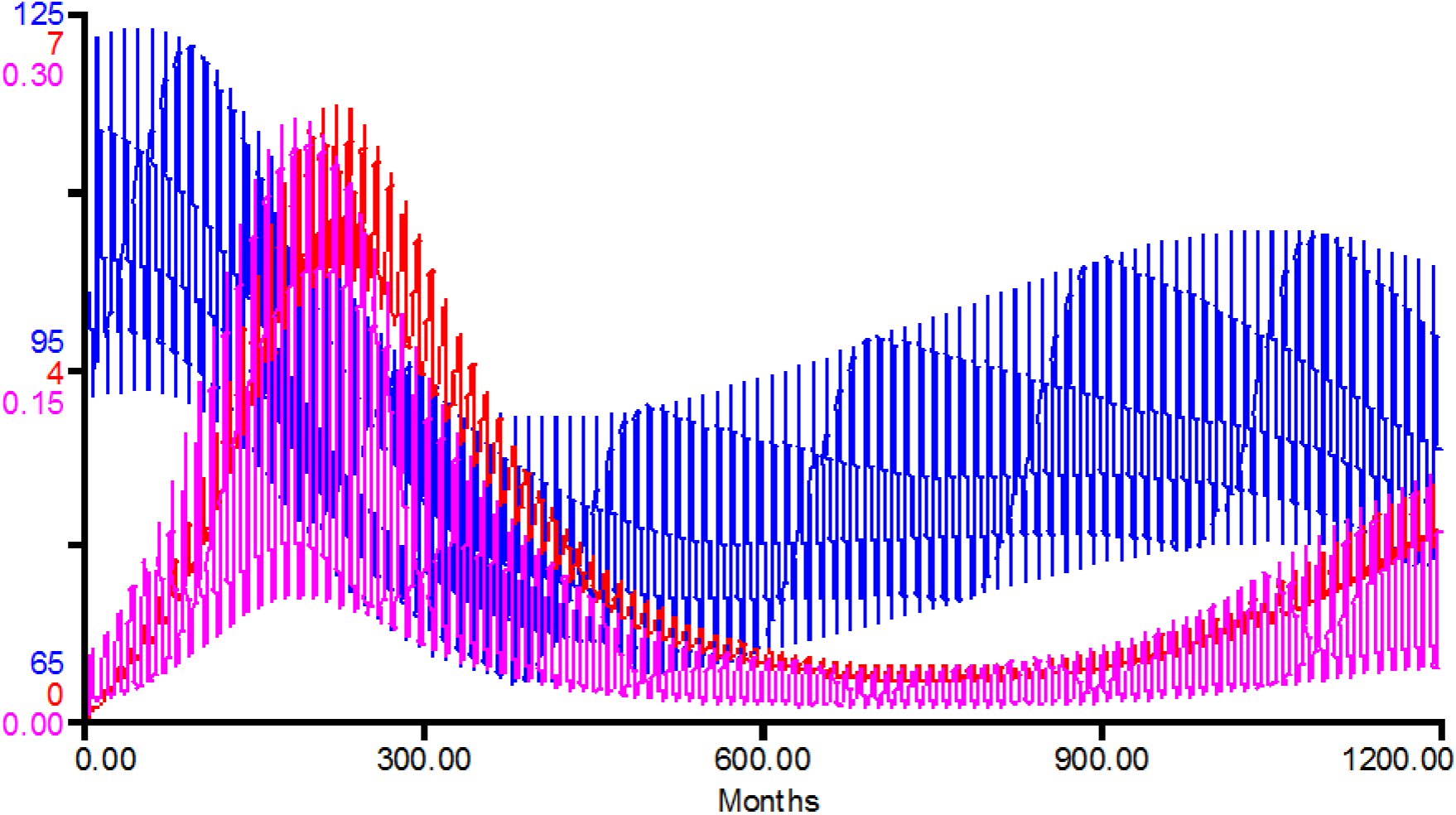
First realistic run of classic SEIR model depicting total population (blue), seroprevalence (red), and prevalence (pink). Where β=0.06, α=0.045, and γ=1.2. Seroprevalence and prevalence are percentage values.

### 3.2. Flexible Model - pre-infectious immunity

The alternate flexible model showed no long-term dynamics and quickly reached a stable state. Seroprevalence quickly increased to around 7-8%, the population remained stable and the percentage of rabid animals was generally less than 0.1% (Figure 2).

**Figure 2.**
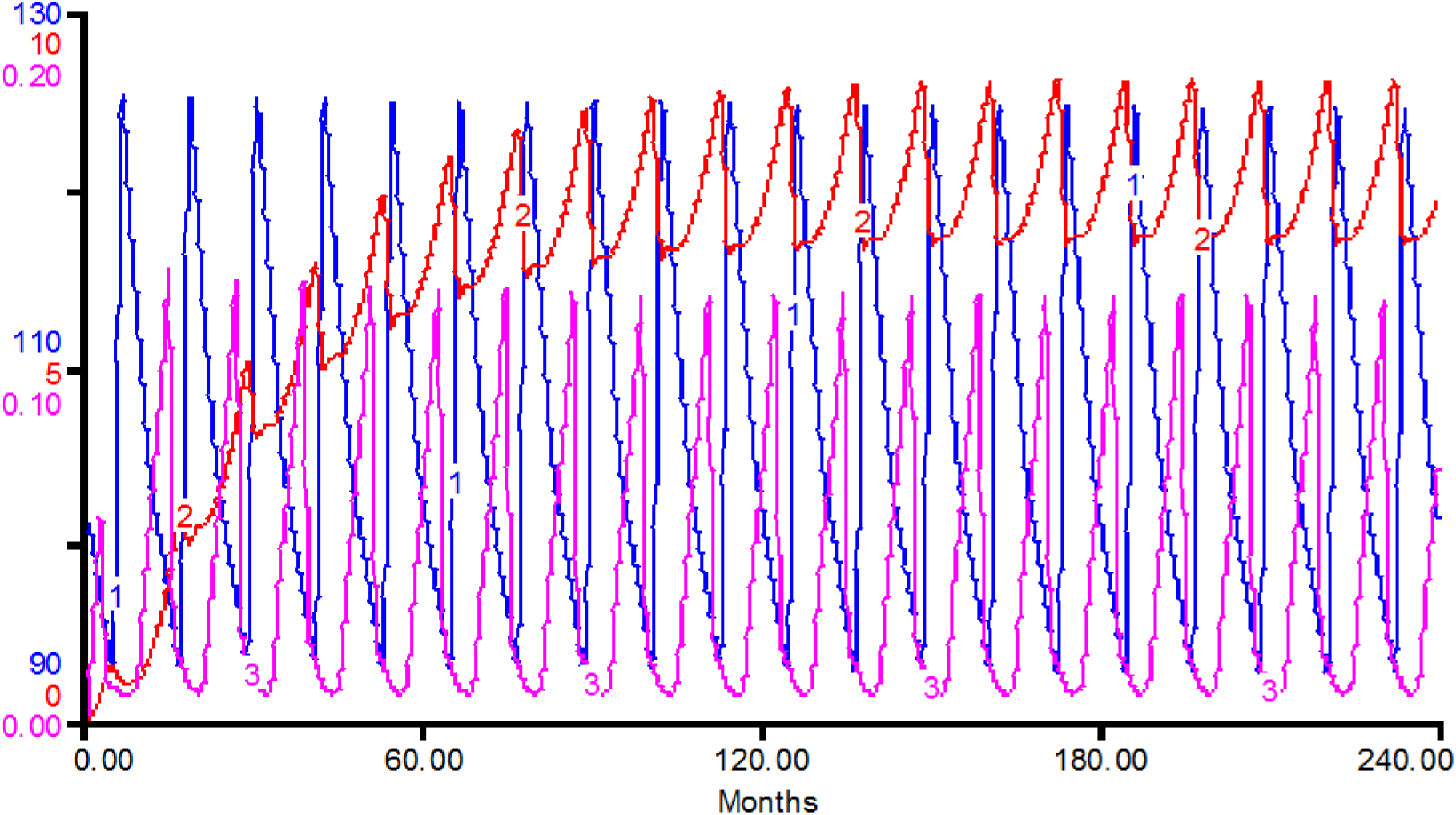
First realistic run of the flexible model depicting total population (blue), seroprevalence (red), and prevalence (pink). Where *β*=0.09, *α*=0.04, and *γ*=0.2. Seroprevalence and prevalence are percentage values.

### 3.3. Sensitivity analysis

For the classical model, alterations in both *β* and *α* led to graphically similar outputs with long-term dynamics. Lower values led to a delay of the seroprevalence peak (at 4%) and higher values created a much earlier peak with larger amplitude (6%). Lower values of *γ* led to an earlier peak and higher values led to the delayed epidemic peak. Alterations in prevalence, measured as percentage of rabid individuals, were minimal for all input parameters ranging from 0.106% (when *β*=0.084, or *α*=0.126, or *γ*=0.107) to a maximum prevalence of 0.255% (when *β, α*, and *γ*=0.255). The other limiting factor, total population, never fell below a value of 67.65; meaning a total population reduction of ~33%.

As above, for the flexible model, alteration of *β* and *α* led to similar graphical outputs. Minimum values (*β*=0.036, *α*=0.012) resulted in diminished seroprevalence with a very low stable level (~1%). Maximum value led to a rapid increase to a stable plateaux (10%). When altering *γ* a minimum value (0.9) brought seroprevalence close to 10%. However, the maximum value (8) was limited not by seroprevalence, but by the low the number of rabid animals. With higher values the rabid population ceased to exist. Alterations in prevalence, measured as percentage of individuals rabid, were much more variable than seen in the classic model. Each parameter gave an average minimum prevalence of 0.05% (*β*=0.073, *α*=0.022, *γ*=0.054) and a maximum average prevalence of 0.17% (*β*=0.161, *α*=0.184, *γ*=0.151). There was also less effect on the total population with it never falling below a value of 87%; a total population reduction of ~13%.

From these upper and lower bounds the global range of the model state space was calculated (Table 4) showing the maximum range of each epidemiological parameter that can result in a realistic output. The values produced showed that the flexible model has an input state space 47 times larger than the classic model (shown graphically in Figures 3, 4, and 5). This means that the total range of input values over which the classic SEIR model produces viable output is 2% of the range of the pre-infectious immunity model.

**Table 4.** Variable parameter minimum and maximum values determined through sensitivity analyses. Global range equates to the volume of the state space for each model and was multiplied by 100 to account for extremely small decimal values.

| Model | B | | $\alpha$ | | $\gamma$ | | Global range |
| --- | --- | --- | --- | --- | --- | --- | --- |
|  | Min. | Max. | Min. | Max. | Min. | Max. |  |
| Post-infectious | 0.042 | 0.06 | 0.0225 | 0.045 | 1.2 | 2.28 | 437.4 |
| Pre-infectious | 0.036 | 0.108 | 0.012 | 0.048 | 0.16 | 8 | 20321.28 |

**Figure 3.**
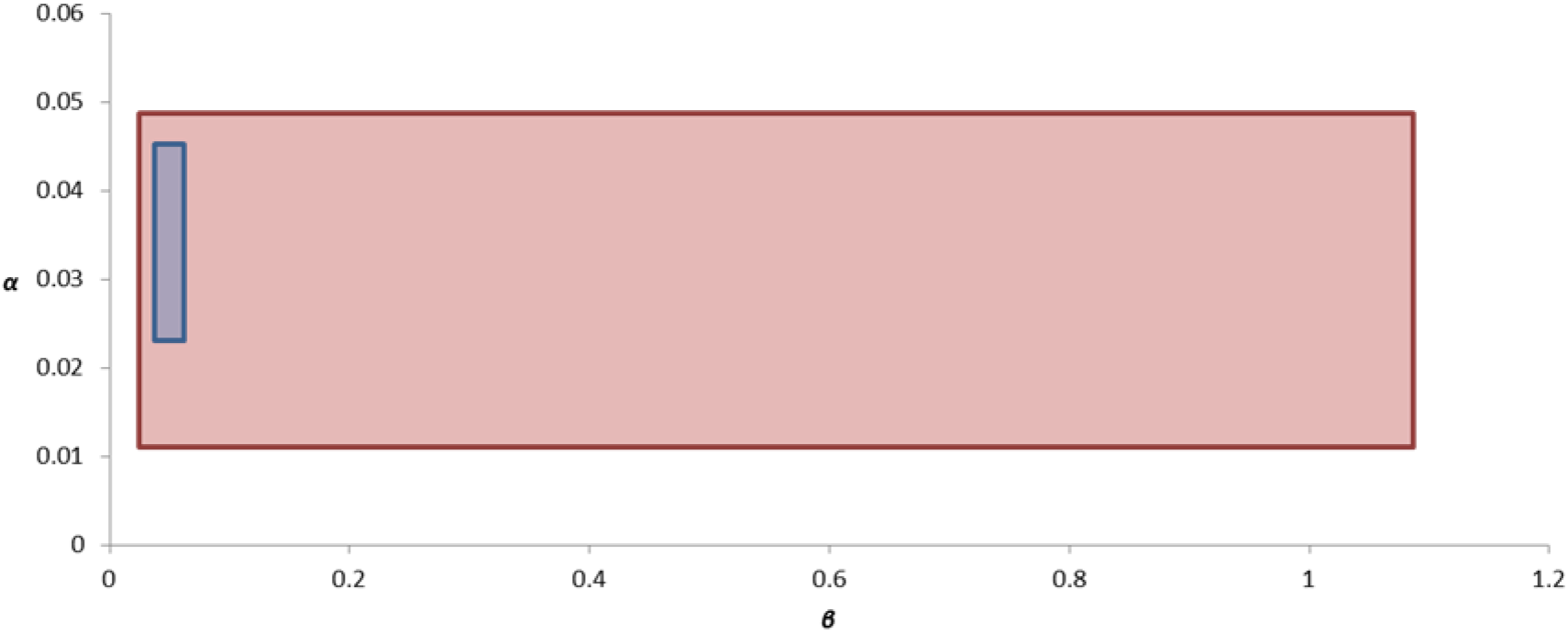
Graphical state space representation for both models; red indicates flexible model, blue indicates classic SEIR model. Plotting maximum and minimum values identified through sensitivity analysis for *α* against *β*.

**Figure 4.**
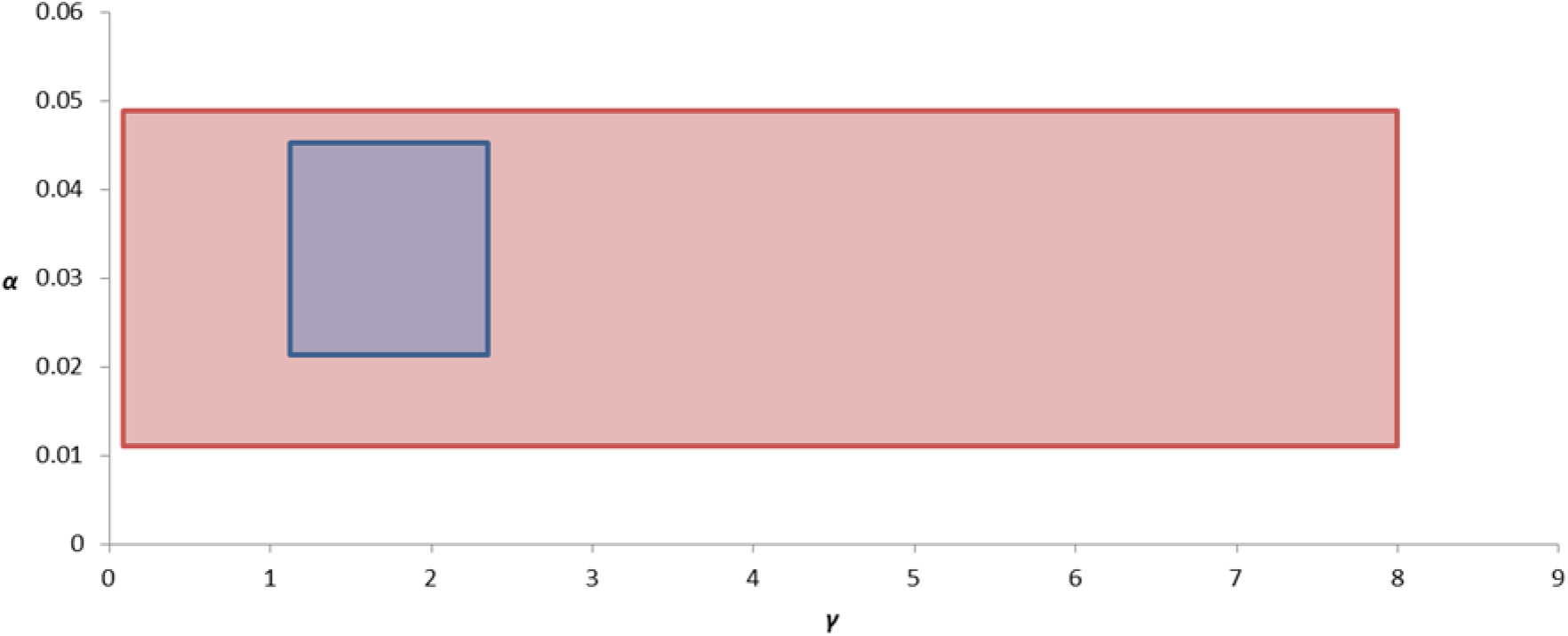
Graphical state space representation for both models; red indicates flexible model, blue indicates classic SEIR model. Plotting maximum and minimum values identified through sensitivity analysis for *α* against *γ*.

**Figure 5.**
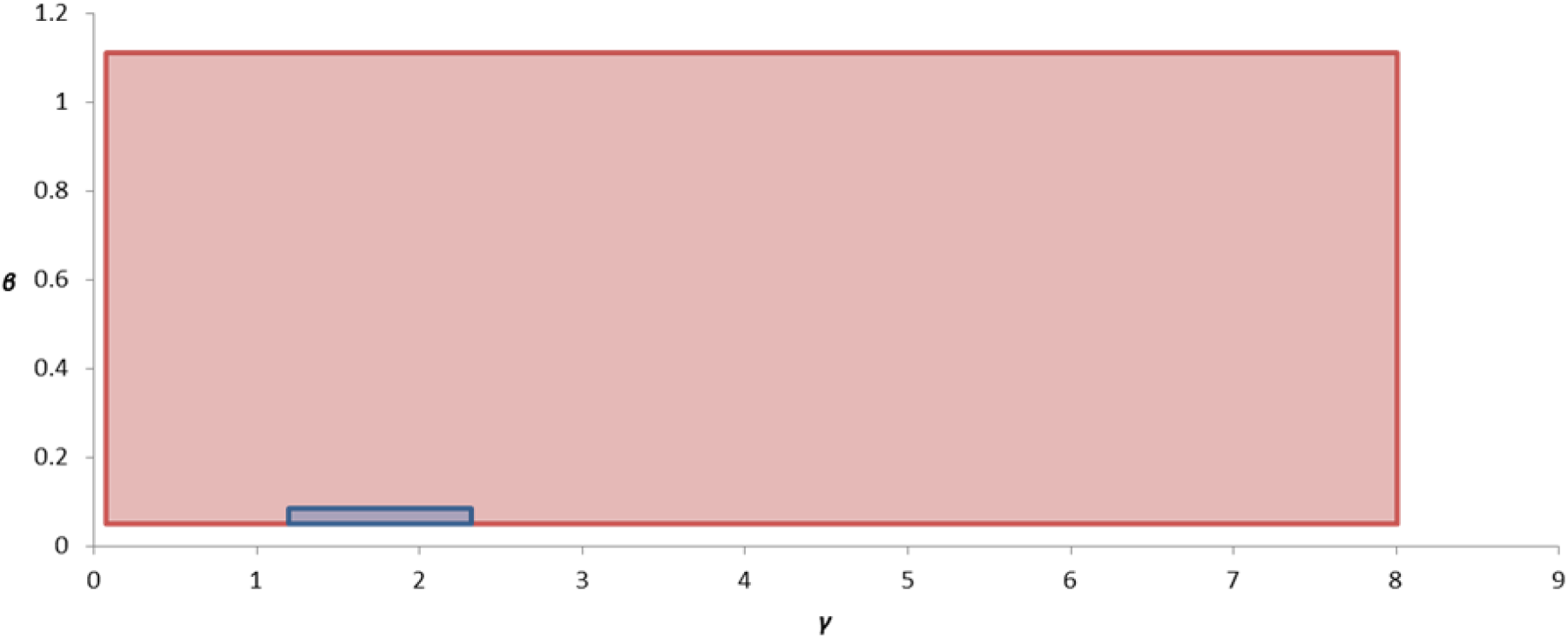
Graphical state space representation for both models; red indicates flexible model, blue indicates classic SEIR model. Plotting maximum and minimum values identified through sensitivity analysis for *β* against *γ*.

## 4. Discussion

Recent epidemic events have led to close scrutiny of bat species as potential vectors and natural reservoirs of multiple zoonotic diseases (Calisher et al., 2006). Whilst many of these serious infectious zoonoses cause fear worldwide, none approach either the epidemiological mortality rate of the Lyssaviruses (Banyard et al., 2011) or the annual number of human deaths they cause. Unfortunately, the understanding of even the basic pathology of lyssaviruses in their wild bat hosts is poor, and empirical field studies struggle with the inherent difficulties of working with wild bats amplified by the problems of working with apparently rare diseases. Here we try to help focus future empirical studies by formally excluding less likely epidemiological scenarios (i.e. putative pathologies) and prioritising field based epidemiological measures (e.g. immunological) associated with more likely scenarios. Models were created for the purpose of identifying the more plausible pathology for determining the mechanisms of EBLV maintenance within a European bat population. Each model followed a basic SEIR design but whilst the classic model constrained immunity in healthy individuals to those surviving disease (post-infectious immunity), the alternate flexible model allowed for ‘incubating’ individuals to follow alternate pathologies and become either rabid, or immune without disease progression (pre-infectious immunity). Our sensitivity analysis demonstrates that the state-space supporting plausible outcomes from our flexible model is 47 times larger than the classic model. Whilst this is not a true probability of the likeliness of either model, the state space analysis identifies that one model has potential for a much greater variation in input parameters and we suggest is more likely to produce endemic disease in a very variable environment.

Bats may have uniquely developed mechanisms to manage lyssavirus infections and acquire immunity, resistance and minimise mortality, which is tangentially supported by various studies empirical and theoretical studies. Field and laboratory based reports indicate non-clinical or sub-symptomatic lyssavirus infections being maintained in bat populations with low levels of disease prevalence (Rønsholt et al., 1998) whilst the few other theoretical studies on other bat-lyssavirus systems have also made similar suggestions (Dimitrov et al., 2007).

The suggested mechanisms within bat immune systems may go some way to helping understand the processes at work (Baker et al., 2013). Here we will accept the suggestion that bat innate immune systems are relatively inactive compared to their rapid and well-resourced adaptive systems which are used as the primary defence against lyssaviruses (Johnson et al., 2008; Serra-Cobo et al., 2002). Key to this is the ability and effectiveness of the adaptive system to control infection in the period between exposure, usually in the peripheral nervous system, and proliferation of the virus once it has reached the central nervous system (CNS). As with other mammals the immune response or potential control of a lyssavirus infection in the bat CNS has been described as limited (Turmelle et al., 2010). Having Virus Neutralising Antibodies (VNA) in a system does not directly indicate whether or not bats must first have a full blown case of rabies or if there is the potential for abortive infection. However, the influence of dose strength on rapidity of infection suggests the potential for a survival threshold value where minimal viral load infection leads to either a reduced response or simply the expression of VNA (Dimitrov et al., 2007).

Further mechanisms that reflect the processes potentially described by the model can be found in the transmission of various rabies strains between bats. In some bat populations oral swabs of apparently healthy individuals have indicated low levels of viral load (Hughes et al., 2006). These low levels of both viral load and success of virus isolation may imply that viral shedding is intermittent in those animals expressing the disease. Low levels of virus passing through direct contact may help explain the presence of abortive infection.

The maintenance of reportable levels of immunity (seropositive bats) in bat populations is unusual for rabies viruses. However, a similar finding has been shown among some humans in the Peruvian Amazon where, presumably through repeated contact with vampire bats, an increase in immune response occurs without full blown rabies (Gilbert et al., 2012). The persistence of immunity is due to the ability of follicular dendritic cells to capture and retain antigen in the form of immune complexes (Baker et al., 2013). This system allows for the persistence of immunity for prolonged periods of time as well as maintaining a memory of immune responses. EBLV-1 antibodies have been found present in some bats over a year after seroconversion (Amengual et al., 2007). This is reflected in both models through the lack of loss of immunity. However, the key to maintaining a disease within these populations is the inability for some individuals to produce VNAs, either due to rapidity of infection or naïve immune system (Freuling et al., 2009). It is also possible that extremely low levels of virus can evade immune system recognition and therefore elicit no response from the preventative systems (Turmelle et al., 2010). Without a preventative system response it is possible that these animals will become rabid. This may well indicate that in situations such as this, bats could potentially become rabid before reaching seropositivity due to low levels of virus replication. The structure of the pre-infectious immunity model allows for some individuals to become immune without ever being infectious as suggested by the classic SEIR model. However, this model still allows for some individuals to become rabid and succumb to the disease. This possibility agrees with the processes suggested here.

Whilst there is field evidence that provides support towards the preferred model, there are also some studies with substantially different population level parameter values. For example, there is a much higher presence of rabies VNA at 65-70% in some bat populations (Burns & Farinacci, 1955; Steece & Altenbach, 1989) and the existence of a high seropositive epidemic peak was only seen in the classic SEIR model. However, these cases of higher seroprevalence were found in classical rabies in North American bats as opposed to EBLV in Europe. There may be some difference in the epidemiology due to the Lyssavirus strain, or it may be due to species differences in behaviour or immunology.

In experimental conditions there is also a potential for animals that had produced VNA to then become rabid after being reintroduced to the virus intramuscularly (Baker & Zhou, 2015). This implies that immunity is not a static state as occurs in SEIR models, but can be over-ridden in certain circumstances. Unfortunately it is not known exactly what causes immunity to fail in these instances and produce symptomatic rabies. Whether it is due to extreme viral load infection or immune system failure (potentially provoked by environmental stressors), this situation has not been accounted for in either model, impacting on the ability of these models to completely reflect real-life processes. However, due to the fact that there is little information known about the circumstances that are involved in this occurrence it could be argued that until there is a greater understanding of the nuances behind individual variation in the immunological responses of wild animals and the factors associated with its failure then it is difficult to include such an outcome.

The models both suggest seasonal variation in seroprevalence and rabid individuals driven by the annual cycles associated with temperate bat biology. These oscillations could influence the interpretation passive surveillance schemes or be exploited by field-based sampling programmes, although the simulated variation is relatively small. For example, the moribund bats submitted to bat hospitals and subsequently confirmed to have died from EBLV-2 often associated with the late summer / autumn (Amengual et al., 2007). Therefore, this may indicate that the current levels of rabies virus identified in the UK are reflective of the peaks of disease prevalence that occur in the warmer months as opposed to the depressed levels found during and slightly post hibernation (Harris et al., 2007). Unfortunately, bats are difficult to study and experimental sample size is generally very low, so ascertaining limited variance in seasonal dynamics would be difficult. From both models it was determined that the incidence of EBLV in the modelled bat populations was between 0.05% and 0.26% which is between 5 and 26 rabid individuals for every 1000 bats so obtaining an empirical measure of the seasonal variance in incidence would be extremely challenging. This is weakly supported by empirical evidence collated across a number of active surveillance programmes (Harris et al., 2009; Schatz et al., 2013) where virus shedding (i.e. an active infection) was found to be very rare (<1 per 2000 bats) which may reflect the model output compounded by the potential behavioural biases reducing the sampling rate in symptomatic bats and the possibility that the period of infectiousness in recovering bats may be brief.

One key area of potential improvement that could be applied is spatial dynamics. The models assume random mixing and therefore no spatial structure. It is possible that temporal-spatial structuring, which can be quite substantial in bat communities, would affect the conclusions of this modelling approach. Both EBLVs have sub-strains that depend on location and latitude (McElhinney et al., 2013) so larger scale spatial structure may not be important for viral maintenance. However, the lack of EBLV1 in the UK and the high genetic diversity of the host serotine bat may imply that small genetically isolated population cannot maintain Lyssaviruses (Smith et al., 2011).

## 5. Acknowledgements

We thank Dr Aileen Mill for support of CB during her MSc project.

